# Investigating the impact of X-rays on decay: X-ray computed tomography as a non-invasive visualisation technique for sediment-based decay experiments

**DOI:** 10.1101/2024.05.13.593831

**Authors:** Iacopo Cavicchini, Paul F. Wilson, Sam Giles, Jake Atterby, Andy S. Jones, Mark A. Williams, Thomas Clements

## Abstract

Decay experiments are ever increasing in complexity to better understand taphonomic processes. However, adding new variables, such as sediment, can create methodological biases, such as artificial anatomical character loss during exhumation. Non-invasive in situ imaging techniques such as X-ray computed tomography (XCT scanning) could mitigate this, but the consequences of exposing carcasses to X-rays are not fully understood, and evidence regarding the impact of X-rays on internal microbial faunas that drive decay is conflicting. Here, we test whether XCT scanning impacts the decay of *Danio rerio* carcasses within a substrate. Our control experiments show that quartz sand sediment physically stabilises the carcass throughout decay and the sequence of anatomical character loss remains constant, however, both the onset and rate of decay of soft tissues are initially accelerated. Our XCT data show that exposure to X-rays does not cause a deviation from the normal sequence of decay, validating XCT as a non-destructive visualisation method for decay experiments. Furthermore, when accompanied with traditional exhumation and dissection, XCT provides decay data with higher accuracy of character analysis than traditional methods, and allows novel quantitative techniques to monitor physical changes in the decaying carcass (e.g., total volume, build-up of gases, collapse of the body cavity etc.). We also underline limitations with the technique, but our experiment acts as an important ‘stepping stone’ for progression toward non-invasive designs of decay experiments.

## Introduction

Decay experiments are a vital tool for understanding taphonomic processes that organisms undergo *postmortem* (Sansom 2014; Briggs and McMahon 2016; Purnell et al. 2018). Experimental designs have become increasingly innovative, investigating the decay of key organismal groups, as well as how decay rates are impacted by specific environmental variables, such as pH, salinity, pressure, and the presence of different sediment types (e.g. Martin et al. 2004; Sansom 2014; Wilson and Butterfield 2014; Gäb et al. 2020; Forel et al. 2021; Clements et al. 2022). Understanding the role of the last of these variables, sediment, is vital for understanding fossilisation, especially of soft tissues. The burial of organisms is a fundamental yet extremely complex aspect of soft tissue preservation, and encompasses many variables that influence decay, including burial speed, burial depth, and mineral availability (e.g. Brett and Baird 1986; Martin, et al. 2004; Olszewski 2004; Wilson and Butterfield 2014; McCoy et al. 2015; Naimark et al. 2018). Experimental work has shown that the mineralogy of the burying substrate influences decay and the physical stabilisation of the carcass, as well as playing a vital role in the preservational processes (e.g. mineralisation and maturation) which allow the geological stabilisation of non-biomineralized soft tissues(Butterfield 1990; Wilson and Butterfield 2014; McMahon et al. 2016).

However, the addition of substrate to a decay experiment means that the carcass is not visible, and the exhumation and/or extraction of carcasses from substrate can introduce bias by damaging the decaying soft tissues. This results in loss of morphological data, especially if the substrate was acting to provide structural integrity to the carcass, and it can potentially skew our ability to accurately record morphological character presence/absence data in decay experiments that model burial – a possibility discussed in the literature (Wilson and Butterfield 2014; Naimark, et al. 2018). Moreover, this bias can be compounded as a decay experiment continues – specimens that are in advanced states of decay, especially when the integrity of the carcass is compromised, are more prone to disarticulation during extraction, decreasing the accuracy of morphological character identification. Previous experiments have employed mechanical solutions to minimise data loss, such as plastic meshes that underlie the carcass to facilitate extraction, or gentle water streams to remove substrate that adhered to decaying carcasses during extraction (Allison 1988; Sansom et al. 2010; Naimark, et al. 2018). Additionally, the requirement for destructive sampling at each sampling point means it is often necessary to decay multiple individual carcasses simultaneously to fulfil sampling requirements for one experiment. This can increase the financial cost, setup time, and physical resources required for representative sampling. Moreover, as the number of carcasses increases, minute biological variation in individuals may introduce minor biases during long experimental runs, leading to sampling point anomalies. It is, therefore, an important pursuit to investigate new methods that could alleviate these experimental issues, and non-destructive imaging techniques like X-ray computed tomography (XCT scanning) have been suggested to replace traditional exhumation and dissection.

Selly and Schiffbauer (2021) were the first to utilise XCT to image a decaying arthropod carcass suspended in plastic beads, demonstrating that high resolution external images can be collected *in situ* in a taphonomic experiment. While this study yielded positive imaging results of the carcass, it highlighted two gaps in our current knowledge. Firstly, it is very difficult to image internal soft tissue structures via XCT scanning without use of a staining agent (Metscher 2009a). Secondly, it is unclear what effects X-rays have on both the rate of decay and sequence of morphological character loss (as per Sansom *et al*. 2010). One of the primary drivers of decay is bacterial metabolism (see Benbow et al. 2019), and the potential impact of exposure to X-rays on the necrobiome of a carcass is unconstrained. As noted by Selly and Schiffbauer (2021), studies investigating the impact of X-rays on bacteria and microbial communities have conflicting findings. High levels of X-ray exposure (much higher than utilised in XCT scanning) are reported as an effective antimicrobial sterilising technique for food items (Mahmoud 2012; Mahmoud et al. 2015), and exposing soil samples to X-rays kills selected bacterial groups (Fischer et al. 2013). Contrary to these findings, other investigations report that X-ray exposure has limited to no impact on soil bacterial communities (Zappala et al. 2013; Schmidt et al. 2015). Should X-rays impact the sequence of decay, then this would render XCT as a non-viable technique for decay experiments. As a counter to the potentially detrimental impact of X-ray exposure on decay-related bacterial communities, Selly and Schiffbauer (2021) suggested that inoculum could be added after each imaging session, however, this could introduce a non-repeatable, confounding variable (see Clements et al. *in prep*).

Here, we report a detailed investigation into the use of XCT scanning as a potential method to non-invasively image a decaying carcass *in situ* within sediment. We undertook a series of decay experiments utilising zebrafish (*Danio rerio*) as our model organism, collecting high-resolution anatomical character presence absence data throughout decay. This study is the first, to our knowledge, to constrain the sequence of anatomical character loss in a vertebrate. Our key finding shows that X-rays have an impact on decay rates but, importantly, do not impact the sequence of character loss, and is a viable technique for decay experiments. Along with these results, our experiment reinforces some limitations of XCT imaging, predominantly the difficulty in resolving the internal anatomy due to lack of contrast between soft tissues. However, this work acts as an important proof of concept, and we outline our ongoing studies focused on designing new methods to overcome these limitations.

## Materials and methods

Zebrafish (*Danio rerio*) are common model organisms in biological studies and detailed anatomical data is available (Menke et al. 2011). Mixed sex, wild-type, AB strain, adult zebrafish (Harper and Lawrence 2016) that had reached the end of their reproductive life were sourced from the husbandry unit at the University of Birmingham. The fish were euthanised using MS222, as per the University of Birmingham’s policy on the use of animals in research, in compliance with the UK Government Guidance on the Operation of Animals (Scientific Procedures) Act, 1986 (Home Office 2014). All fish were used immediately after euthanising, on day 0 of the experiments outlined below.

### Experimental control (CTRL) and sediment matrix experiment (SNDCTRL)

Individual *D. rerio* (*n* = 96) were placed in 200 mL plastic containers filled with 100 mL of deionised water. For ease of extraction, the carcasses were placed on top of a circular piece of plastic mesh, following Sansom *et al*. (2010). All jars were kept at a constant temperature of 25°C using an IPS260 Memmert maturation chamber for the duration of the experiment. For the control experiment (CTRL), *D. rerio* carcasses (*n* = 48) and the plastic mesh were placed in deionised water and then sealed for the duration of decay. For the second experiment (SNDCTRL), *D. rerio* carcasses (*n* = 48) and the plastic mesh were buried in approximately 100 grams of 0.1 - 0.5 mm quartz sand (VWR International) and then sealed.

A sampling strategy was designed to identify the sequence of morphological character loss, with a more intensive sampling frequency at the initiation of the experiment. Capturing the early phases of decay at high resolution was necessary, as previous experiments have shown that decay of internal organs is rapid (e.g. Sansom *et al*. 2010; Clements *et al*. 2022). The sampling points were as follows: 1 fish on day 0; 4 fish on days 1-4; 2 fish on days 5-7; and 1 fish on days 8-14, 16, 18, 20, 22, 24, 26, 28, 30, 33, 36, 39, 42, 45, 50, 55, 60, 65, 71, 78, 85, 92, and 120. Each carcass was carefully extracted and dissected under an optical microscope.

40 morphological characters were assessed in each carcass (see Appendix S1), recording if they were in life position, and assigning one of four decay states: pristine (visually same condition as immediately *postmortem*), decaying (morphology altered but identifiable), onset of loss (morphology altered and difficult to identify), complete loss (no longer observable or recognisable) as per the terminology designed by Sansom *et al*. (2010), with one additional character state (onset of loss) to increase the resolution of our dataset. We used 1.5 IQR tests to check for outliers in the datasets of both experiments.

In order to quantify the differences between the sequence of anatomical character loss of the CTRL and SNDCTRL experiment we assigned a numerical score to the codings of each experiment following the revised methodology of Sansom *et al*. (2010): “pristine” codings received a score of 0, “decaying” and “onset of loss” were scored as 0.5 and “complete loss” was scored as 1. The total score was calculated for each specimen, allowing a decay-through-time analysis, and Spearman’s rank correlation coefficient was used to investigate variations in the sequence of anatomical character loss between specimens.

### Computed tomography experiment

To test the impact of X-rays on decaying carcasses, we utilised an experimental protocol similar to the previously described SNDCTRL experiment: *D. rerio* carcasses were individually placed in separate 200 mL plastic containers, filled with 100 mL of deionised water and then buried in approximately 100 grams of medium to fine quartz sand. One of the requirements of XCT scanning is that X-rays must penetrate the entire sample from multiple directions, requiring that either the sample or the X-ray source and detector rotate during imaging. This study utilises the former method, and this presents an issue when designing decay experiments, as sample movement can cause carcass agitation, known to cause disarticulation of the organism (Briggs and Kear 1993), as well as preventing accurate images to be obtained. In the past, this issue was countered by utilising plastic beads to stabilise a shrimp carcass during XCT scanning (Selly and Schiffbauer 2021). We use a quartz sand medium to stabilise the zebrafish carcasses allowing them to be imaged in situ, however, this meant that it was then necessary to test the impact of sediment on the sequence and rate of decay.

, as well as preventing accurate images to be obtained. Selly and Schiffbauer (2021) countered this issue by utilising plastic beads to stabilise a shrimp carcass during XCT scanning. Here, the quartz sand medium stabilises the carcasses, allowing them to be imaged *in situ*.

The X-ray samples were split into two batches: one batch was scanned at the start (day 0) and end (day 50) of the experiment (*n* = 4, hereafter referred to as samples A, B, C, D), while the second batch was only scanned at the end of the experiment (day 50; *n* = 4, hereafter coded E, F, G, H). At the termination of both experiments, the fish carcasses were extracted from the sediment and dissected. Between scans, the jars were kept in a water bath in a temperature controlled environment (18-22°C) in proximity to the XCT machine. Although this setting was not ideal, as temperature was not as regulated as in the control experiment, it was necessary in order to keep the carcasses as close to the XCT machine as possible to reduce agitation between and during the scanning sessions.

XCT scanning was carried out using a Tescan Unitom XL at the National X-Ray Computed Tomography (NXCT) facility at the Centre for Imaging, Metrology and Additive Technologies (CiMAT) at the University of Warwick. All scans used the following settings: 140 kV energy, 45 W current, 1 mm tin (Sn) filter, two frames per projection, exposure time 370ms, effective voxel size of 45μm. The acquired image stacks were prepared into tomographic datasets using ImageJ software (v. 1.53t), manually cropping empty space around the fish carcasses. Segmentation of the 12 3D models was undertaken using Mimics Materialise software (v. 20 and 22; http://biomedical.materialise.com/mimics; Materialise, Leuven, Belgium), relying mostly on density threshold tools and automated algorithms, limiting the amount of hand-drawn segmentation to a minimum. This method prevents potential human bias that can be introduced during segmentation, at the cost of a grainy surface texture.

To quantify the differences between SNDCTRL and the specimens that were scanned using XCT (samples A-H), we used binary coding (presence/absence) for all specimens involved and omitted the “gonads” character, as this can be hard to observe in male zebrafish. To compare the decay rates, we calculated the percentage of absent characters for A-H and for specimen # 39 from SNDCTRL (this specimen is the corresponding carcass that was extracted at day 50). To test whether our ability to accurately code characters in a decaying carcass is impacted by the method of data collection, we compared the percentages of presence/absence from two sources: (1) character presence/absence solely from the dissections performed after XCT scanning and (2) character presence/absence from the dissections and incorporating the characters observed via tomograms.

Finally, to test whether the sequence of anatomical character loss is impacted by XCT scanning, we ranked the characters of A-H and specimen # 39 from SNDCTRL using the revised methodology from Sansom *et al*. (2010, see previous paragraph) to perform Spearman’s rank correlation coefficient: firstly, scoring the characters of A-H only from dissections, then only from XCT images, and finally coding characters that were recognisable with either method.

### 3D model mesh comparison

Specimens A, B, C and D were XCT scanned twice (on day 0 and day 50), allowing 3D volumetric mesh models for each specimen at the start and end of the experiment to be created. We imported the mesh pairs into CloudCompare (Ver. 2.13, 2023), then aligned them using the CloudCompare Fine Registration (ICP) algorithm allowing the geometry (and thus volume) of the two meshes to be compared. We used the Cloud to Mesh distance algorithm to compute the distance in millimetres between each point on the surface of the day 50 mesh and the closest point on the surface of the day 0 mesh; the difference was output as a heat map plotted onto the day 0 meshes.

## Results

### A chronological report of the control experiment: D. rerio carcasses decaying in water (CTRL)

The first visible signs of decay observed in the zebrafish carcasses occur on day 1 (Fig 1; Fig. S1-S2). The eye becomes clouded, darker, and often a small air bubble forms below the lens, while the heart becomes smaller and loses its normal red coloration, becoming pink. On day 2 the female gonads begin to lose structural integrity – this paired organ contains hundreds of oocytes and upon rupture, the oocytes can fill the body cavity; if this occurs, they remain observable for the duration of the experiment. Between day 2 and 6, the internal organs start to show signs of decay: the liver and pancreas, normally visible around the intestinal bulb as areas of (respectively) dark red-brown and yellow-green coloration, become harder to identify and start to lose structural integrity (day 2); the three parts of the intestine (anterior, bulb, and posterior) lose their original tube-like shape and appear flattened (day 2); the gills start losing colour, transitioning from red to pink (day 3); and both swim bladders rupture (day 3-4). The zebrafish kidney is detectable as a filamentous structure attached to the roof of the coelom, extending from the back of the skull almost to the end of the body cavity. It remains visually identifiable throughout the whole experiment, but parts of it disarticulate and detach from the coelom roof from day 4. The external pigmented stripes on the skin and fins begin to fade from day 4 as well, although they are never entirely lost. The skin becomes flaccid and more difficult to pierce between day 4 and 5, but rarely splits or ruptures. From day 8, the zebrafish bones, typically starting with the skull and pectoral girdle, become translucent and disarticulate readily during dissections. The heart, liver, and pancreas are often completely absent by day 11. On day 11, the brain “liquifies”, completely losing its structural integrity, with the ‘liquid’ remaining confined within the braincase. From day 13, the fin rays lose their normal rigidity and articulation, and the membranes no longer sustain the structure: as a result, the fins are found compressed and bent. The anterior swim bladder is undetectable after day 13, except on a few occasions where the deflated membrane persists. On day 16, the gill filaments start to lose ultrastructure, becoming gelatinous.

**Figure 1:**
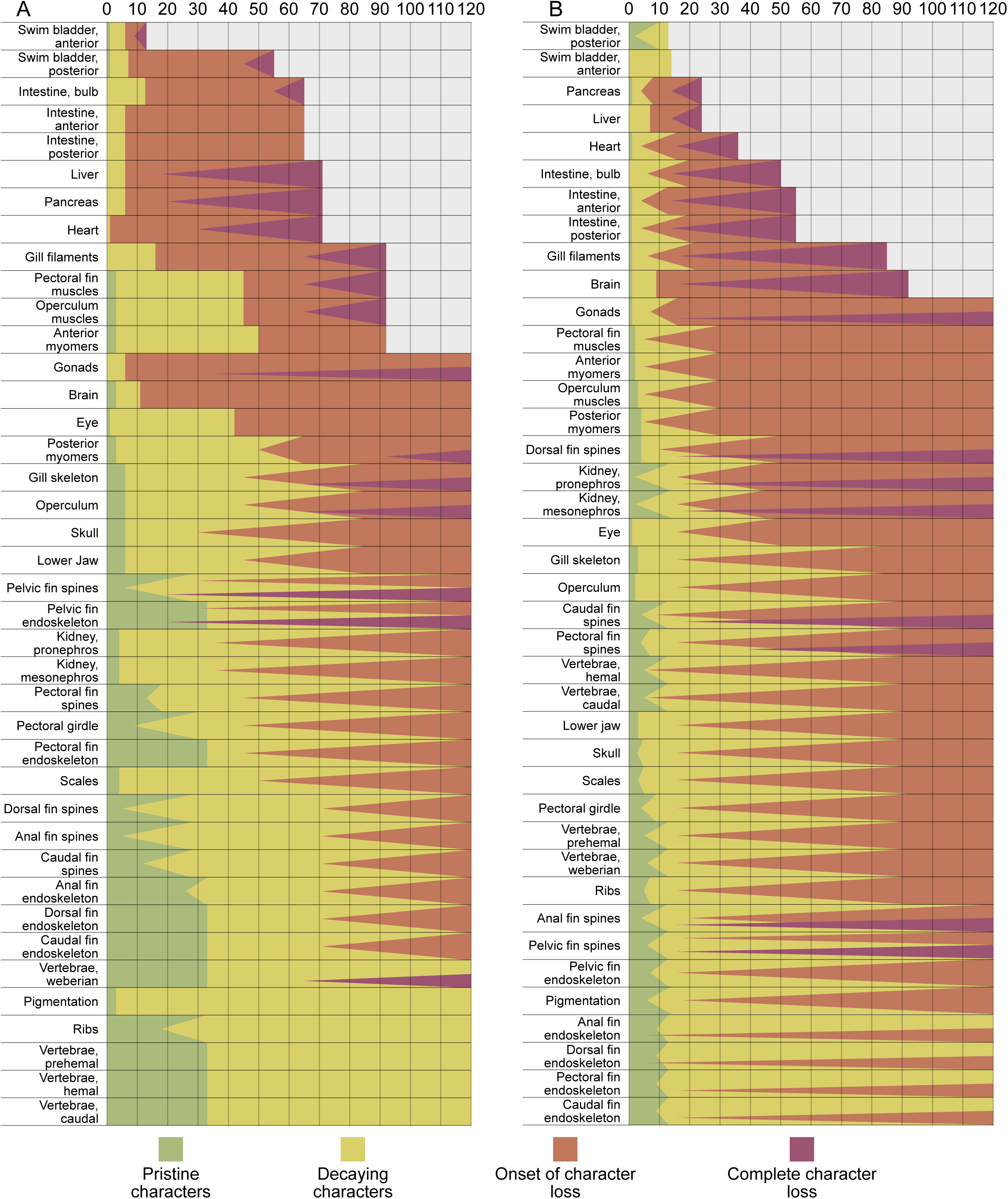
Sequence of anatomical character loss in zebrafish decaying in water (CTRL; A) and quartz sand substrate (SNDCTRL; B). Experiment days indicated at top and key of character states at bottom. Characters are ordered based on the last occurrence during the experiment (most decay resistance at bottom). Character score ties were broken based on (in order of priority): transition to onset of loss; first occurrence of complete loss; first occurrence of onset of loss; transition to decaying, first occurrence of decaying. Triangles indicate transition periods between character states. Character state definitions: pristine (same condition as at *postmortem*), decaying (morphology altered but easily identifiable), onset of loss (morphology altered and difficult to identify), complete loss (no longer observable or recognisable).

By day 30, the intestine loses structural integrity and can rupture. On day 36, the male gonads decay and are no longer identifiable. At this time, the vertebral column and ribs become translucent and prone to disarticulation. Around day 40, the skull, pectoral girdle, and scales become semi-transparent and disarticulate more readily during dissections. Fin rays are often found with the tips missing and lose all remaining rigidity. The fin membranes lose rigidity as well and become saggy, but usually remain articulated. The eyes are typically opaque and, if the cornea loses structural integrity and collapses, the eye looks completely black. At this time, the muscle tissues lose their structure and become gelatinous.

On day 45, the posterior swim bladder decays and is not detectable anymore, followed on day 65 by all the parts of the intestine. The heart, liver, and pancreas decay by day 71 and muscle tissues and gill filaments become unidentifiable on day 92.

The water in the plastic jars becomes progressively murkier between days 2 and 10, with an increasing amount of floating residue. Between day 13 and day 45, a film of dark hydrolysed fats (adipocere) often forms on the water surface. By day 30, any suspended residue falls from suspension, settling on the jar floor, and the water becomes clear again. An exception to this occurs if portions of the carcass become disarticulated, causing the water to remain murky and filled with suspended residue. The residue was not identified during this investigation.

During the control experiment, most fish carcasses conformed to the ‘normal’ sequence of anatomical character loss described above, but some carcasses deviated from it. In the second carcass sampled on day 6, elements of the head and the pectoral girdle were disarticulated from the rest of the body and unidentifiable, which caused early loss of the heart and gill filaments (Fig. 2; Fig. S1-2). The carcass sampled on day 24 was in an advanced decay state compared to the other individuals sampled at this time point, with complete loss of the head, pelvic fins, and all internal organs (except for the kidney).

**Figure 2:**
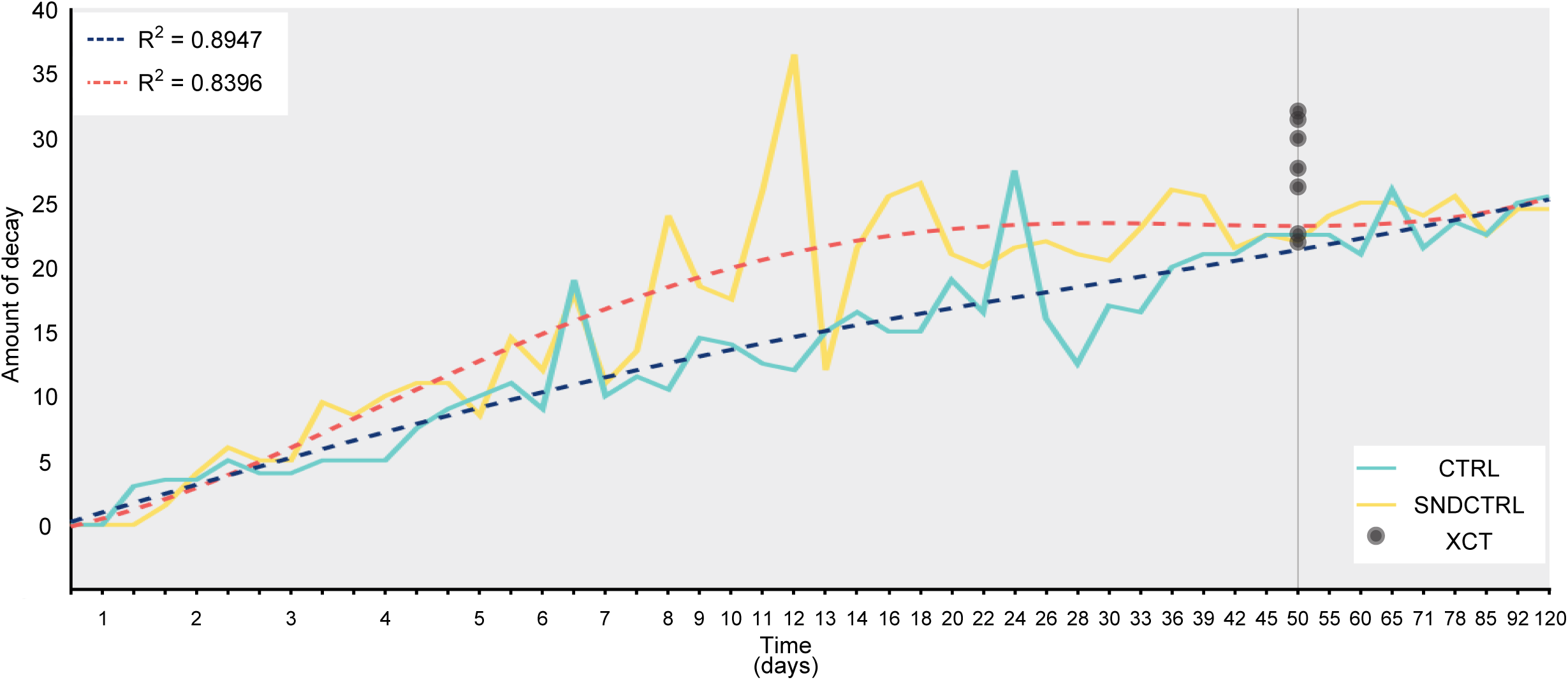
Decay rates of zebrafish in water (CTRL) and in a quartz sand substrate (SNDCTRL). Cyan line: zebrafish in water (CTRL experiment); blue dotted line: line of best fit for CTRL. Yellow line: zebrafish in quartz sand and water (SNDCTRL experiment); red dotted line: line of best fit for SNDCTRL. Vertical axis: amount of decay is accumulative character state score (for each character: pristine: 0; decaying, onset of loss: 0.5; complete loss: 1. Minimum: 0 (pristine fish); maximum: 40 (complete loss of the whole carcass).

### A chronological report of carcasses decaying within a sand matrix (SNDCTRL)

The first visible signs of decay occurred on day 2 (Fig. 1; S1-S2). The eye appears glassy, opaque and develops a small air bubble below the lens, the anterior swim bladder and female gonads rupture, the gill filaments and the brain begin to lose integrity, the heart appears smaller and darker, the liver and pancreas start to lose their overall structure but remain detectable as does the the intestinal bulb, and each part of the intestine appears darker and reduced in volume. The posterior swim bladder persists longer than the anterior one, retaining its structure and the gas inside it as late as day 10. Between days 3 and 4, the skull and pectoral girdle bones become translucent and less resistant to disarticulation during dissections. Between days 4 and 5, the fin rays lose rigidity, and the fins can be found bent in unnatural positions. In the following days, the loss of volume of the heart continues (day 4-10), the liver and pancreas lose their overall structure (day 7-10), and the three parts of the intestine (anterior, bulb and posterior) appear flattened (day 4-7). From day 9, the vertebral column becomes less resistant to disarticulation during dissection, and individual vertebrae become translucent. The brain finally loses any detectable structure on day 9, although the liquified remains normally persist, trapped inside the braincase. On day 14, both swim bladders become undetectable and the muscles become gelatinous. From day 16, portions of the filamentous structure of the kidney can detach from the roof of the coelom. From day 20, the fin rays often lose their distalmost portions and the scales become translucent and hard to detect. The liver and pancreas become unidentifiable on day 24, while the intestine remains detectable albeit flattened, discoloured, and often with its parts disconnected from each other. On day 33 the skull, pectoral girdle, and axial skeleton become transparent and have almost no resistance to disarticulation during dissections. On day 36, the heart becomes unidentifiable. The same happens to the gill filaments on day 50, while the entire intestine and the brain become unidentifiable on day 55.

The water in the jars becomes murky from day 2, and the suspended organic residue persists until day 20, when it settles on the surface of the sand. From day 7, an adipocere layer develops on the water surface: it can vary in colour from grey to black and persists for the whole experiment.

A number of carcasses deviated from the typical sequence of anatomical character loss (Fig. 2; Fig. S1-S2) during the experiment. In the specimen sampled on day 8 the gill filaments, heart, liver, pancreas, swim bladders and kidney were already unidentifiable, its skeleton was translucent with little resistance to disarticulation and the fin ray tips were partially missing. Another carcass sampled on day 11 had the body cavity filled with unrecognisable remains of the internal organs and the ovaries, its fin rays had the tips missing, the anal fin was not recovered, and its axial muscles already had a gelatinous texture. The carcass sampled on day 12 was substantially decomposed and disarticulated – the entire body was almost completely lost, save for a few scales, the posterior-most portion of the trunk with associated vertebrae and musculature, and fragments of the jaw.

In contrast to specimens that decayed quicker than expected, the day 13 specimen deviated from the expected sequence of anatomical character loss by decaying more slowly. The fin rays showed only minor signs of decay, the vertebral column was in pristine condition and all the internal organs were still recognisable, although the brain had lost its structural integrity.

The carcasses sampled on days 16 and 18 showed decay that deviated from the norm, with notable differences in the skeletal elements. In both fish, the fins were either unidentifiable or in advanced decay states, with no rigidity and the tips of the fin rays missing. Furthermore, in both specimens, the vertebrae, ribs and skull were transparent and opposed no resistance to disarticulation during dissection. In addition, in the day 16 specimen the posterior portion of the vertebral column and the caudal fin were disarticulated from the rest of the carcass, while in the day 18 specimen the head was disarticulated from the rest of the body. Finally, the fish sampled on day 85 represents another occurrence of slower decay rate. The skeleton was almost intact, retaining some resistance to disarticulation, and some delicate internal organs persisted – the heart, gill filaments and brain were greatly reduced in volume and had lost most of their structural integrity, but were detectable. The fin rays lost their rigidity but the overall structure was intact.

### Comparison between the control experiment (CTRL) and carcasses decaying in sediment (SNDCTRL)

We compared the numerical scores of the carcasses in the CTRL and SNDCTRL experiments, obtained following the revised methodology of Sansom *et al*. (2010) (see Methods section). The scores of carcasses in the SNDCTRL are typically higher than those of the CTRL between day 2 and day 85, indicating a higher amount of decay, and also have a larger range than the CTRL (Fig. 2). The maximum difference is reached between day 8 and day 18, and subsequently the two experiments converge to similar amounts of decay for the last 3 sampling points (days 85-120). The rate of decay for the CTRL is constant throughout the experiment (Fig. 2). In contrast, the initial rate of decay for the SNDCTRL is faster than the CTRL until day 12, when the rate begins to slow, and then subsequently plateaus around day 16 - 20. Eventually, around day 85 the rates of decay of the CTRL and SNDCTRL intersect. Detailed investigation of the timing of anatomical character loss using the four state character coding (pristine, decaying, onset of loss, and complete loss) demonstrates that burial within a substrate impacts, not only the rate, but also the onset of decay.

Anatomical characters typically lose their ‘pristine’ coding faster in the SNDCTRL than in the CTRL samples (Fig. 1; Fig. S1-S2). We see a similar trend in that the percentage of anatomical characters coded as ‘onset of loss’ also increases faster in the SNDCTRL specimens than those in the CTRL (Fig. S2). There are two obvious spikes in the proportion of lost characters: one on day 24 for the CTRL and one on day 12 for the SNDCTRL (Fig. 2; Fig. S2). A 1.5 IQR test reports that both specimens are anomalous and are statistically outliers in their respective datasets. When the numerical scores are used to assign a rank to each character (thus quantifying the sequence of character loss), the Spearman’s rank correlation coefficient and the associated *p* value indicate that there is no significant difference between the sequence of anatomical character loss between the fish decayed in the CTRL and in the SNDCTRL (R_S_ coefficient = 0.87, *P < 0.0001*; Table S1). In both experiments the swim bladders are the first to be lost, followed by the rest of the organs in the body cavity (liver, pancreas, heart, intestine) and the gill filaments. Kidney, brain, gonads and musculature can be lost or persist for the duration of the experiment, while the skeleton is always persistent at the timescale of our investigation (Fig. 1).

### Cataloguing the morphological decay of carcasses using XCT scanning

XCT imaging allowed us to obtain 12 renderings of zebrafish carcasses *in situ* (Fig. 3). The resulting 3D models have a rough surface texture due to sand grains adhering to the external surface of the carcasses, which segmentation algorithms fail to separate from the carcass. This problem was mitigated as much as possible with the available automated algorithms, because they were found to be more precise and consistent than manual segmentation. Despite this, the resolution of the scans is high enough that details of external features such as the eye, mouth, operculum, and fins can be identified from the 3D models (Fig. 3A1-H). All of the carcasses remained intact for the duration of the experiment, with limited visible disarticulation. At the initiation of the experiment, the carcasses have gas trapped in the body, either within the still intact swim bladders or as isolated bubbles within the intestinal tract or the mouth. As the carcasses decay the swim bladders rupture, and the gas they contain is released into the coelom. Bubbles of gas are still visible in all of the day 50 XCT images, and the gases escaped the body and the sediment cover during extraction. There were two exceptions to this trend: specimen G, XCT scanned on day 50 only, was devoid of gas; and specimen C, which may have suffered accidental damage to the swim bladders during experimental set-up, and has a minimal volume of gas visible in the day 0 XCT scan (Supp. Mat.). The day 50 scan of specimen C, however, shows more and larger gas bubbles than the day 0 scan, probably as a result of gas build up during decay.

**Figure 3:**
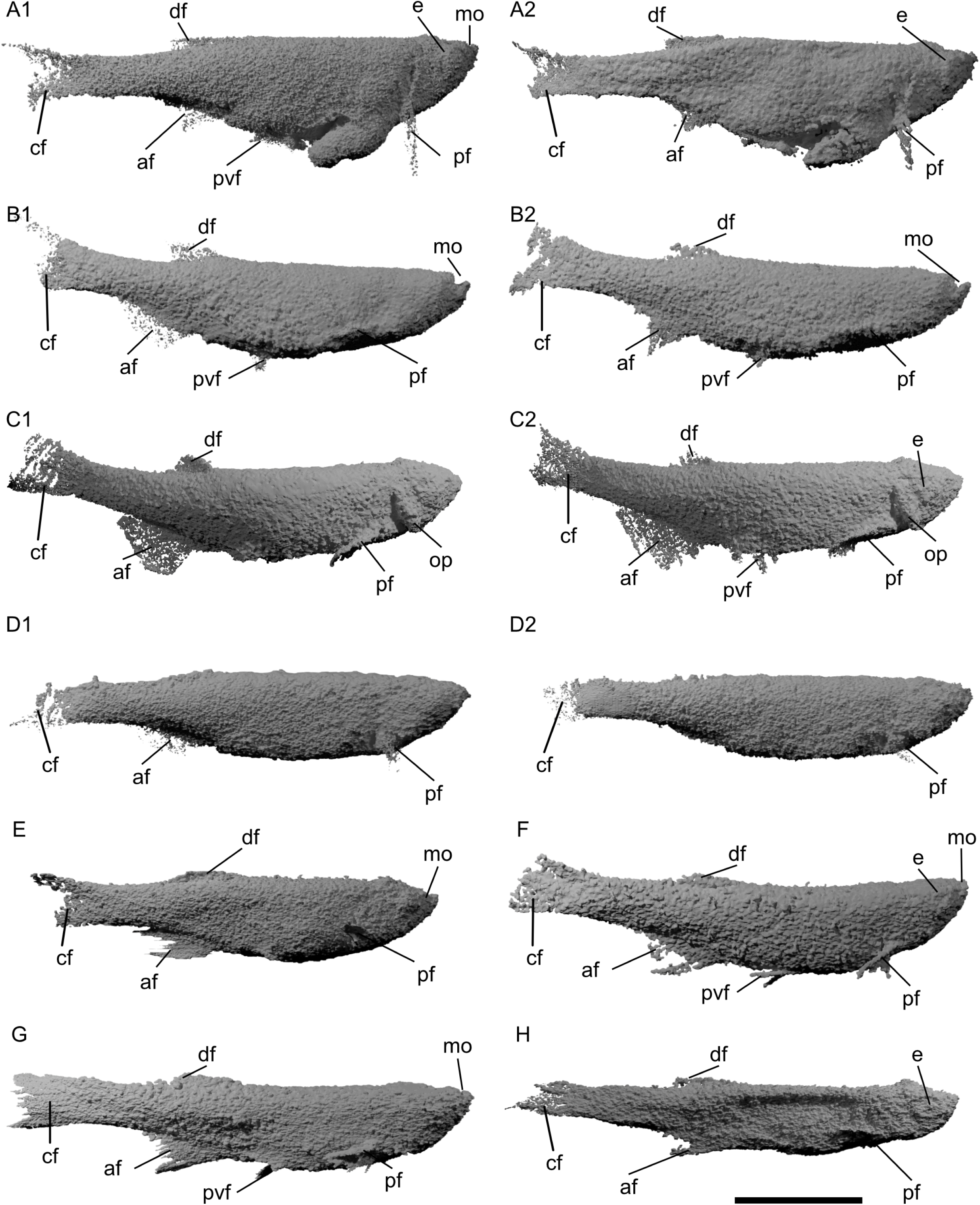
3D XCT models of zebrafish in various stages of decay in right-lateral view. A1-A2, specimen A on day 0 and on day 50; B1-B2, specimen B on day 0 and day 50; C1-C2, specimen C on day 0 and day 50; D1-D2, specimen D on day 0 and day 50; E, specimen E on day 50; F, specimen F on day 50; G, specimen G on day 50; H, specimen H on day 50. Scale bar: 1 cm. Abbreviations: af, anal fin; cf, caudal fin; df, dorsal fin; e, eye; mo, mouth; op, operculum; pf, pectoral fin; pvf, pelvic fin.

The scans also allow us to observe the orientation and shape of the carcasses. Most specimens were oriented laterally (*n* = 4), or dorsolaterally (*n* = 3), while only one specimen was oriented dorsoventrally. Specimen H was laterally flattened – more so than the other specimens (Fig. 3H). This suggests collapse of the coelom and loss of the internal organs, which was confirmed by dissecting the carcass (see below). Specimen A had a rupture along the ventral side, probably developed during burial, with a portion of the intestine extruding outside of the body cavity and detectable in the XCT images on day 0 and day 50 (Fig. 3A1-A2).

All carcasses in the XCT experiment were dissected immediately after they were scanned on day 50 in order to allow data on the state of morphological decay to be collected and compared with data from the SNDCTRL (*n* = 8). Here, we report the state of morphological decay of the scanned carcasses at day 50.

*Specimen A*: the body had disarticulated into several unidentifiable fragments and some distinct sections of articulated spine and associated axial musculature. The head had disarticulated from the rest of the body and the eyes were darkened, with both lenses unidentifiable. Some larger fragments of the body retained the pigmented stripes on the skin. No fins were recovered. No internal organs were identifiable. The skull and the discernible portions of the vertebral column were semi-transparent and disarticulated readily when examined during the dissection.

*Specimen B*: the body had disarticulated into several pieces, the larger of which retained associated skeletal elements and musculature. The head was separated from the body. The eyes were black, but the lens was still intact and identifiable. Only portions of the pelvic and pectoral fins were able to be identified. The brain and the internal organs were not identifiable. All parts of the skull and the identifiable portions of the axial skeleton were semi-transparent.

*Specimen C*: the body remained intact. The eyes were dark but the lens was visible and in life position. Most of the musculature was gelatinous but still identifiable. Pigmentation on the body was fading but still visible. The pelvic fins were not recovered, but all others were present. The anal and caudal fins were particularly intact, although the overall structure was starting to lose robustness. The internal organs and the brain were no longer observable. The skull and pectoral girdle were semi-transparent and disarticulated readily during the dissection, while the vertebral column showed a higher resistance to disarticulation during dissection.

*Specimen D*: the body remained intact. The eyes were dark but with the lens intact and in life position. All the musculature was gelatinous but still detectable. Pigmented stripes on the body were faint but visible. All the fins were present, although their robustness was lost. The brain had no structural integrity, however, the liquified remains were still within the braincase. The heart, liver, pancreas, and anterior swim bladder were not identifiable, although the deflated posterior swim bladder was detected. Likewise, the three major sections of the intestine were identifiable, albeit flattened and discoloured. Elements of the kidney were also identifiable on the roof of the coelom. All the skeleton was translucent, but retained articulation, resisting disarticulation during dissection.

*Specimen E*: the body remained intact. The eyes were black but structurally intact, with the lens still in life position. All the muscles were gelatinous but identifiable. Colouration on the body was faded but still visible. The fins were identifiable but had lost their structural robustness. As with the previous specimen, the brain had liquified, but only part of it was still visible in the braincase. The heart, liver, pancreas, and anterior swim bladder were not identifiable. The posterior swim bladder was still present but deflated. The intestine was identifiable but discoloured and flattened. The kidney had disarticulated but fragments were identifiable. The entire skeleton was translucent and retained some resistance to disarticulation during dissections, except for the bones around the skull roof. These bones were transparent and less resistant to disarticulation during dissection.

*Specimen F*: the specimen had disarticulated. Most of the head, pectoral girdle and anterior musculature were not recovered. The eyes were black and the lenses had disarticulated. The posterior portion of the body had disarticulated and most of the associated axial musculature was unidentifiable. Some of the pigmented stripes were visible on the fins and where the dermis persisted. The pectoral and anal fin were not found, and other fins had lost some rays. The fin membranes had lost their robustness. The brain and the internal organs were not recovered. The portions of the skeleton that were identifiable were semi-transparent and disarticulated readily when dissected.

*Specimen G*: the body remained intact. The eyes were black and the lens was still identifiable and in life position. The musculature was gelatinous. The pigmented stripes on the body were faint but visible. All the fins were found albeit without any remaining structural robustness. As with previous specimens, the brain had liquified and a pale matter in suspension (with a similar consistency to milk) was still partially visible in the braincase. The heart, liver, pancreas, anterior swim bladder, and posterior portion of the intestine were not identifiable. The posterior swim bladder was identifiable but deflated. The remaining two parts of the intestine were pale and flattened. Elements of the kidney were still visible. All the skeleton was translucent and retained some resistance to disarticulation during dissection.

Specimen H: the specimen was disarticulated. The head and the pectoral girdle were detached from the rest of the body and the body cavity was ruptured ventrally. The eyes were still visible but entirely black and the lenses could not be identified. The muscles were gelatinous but detectable, including the pectoral muscles. The dark stripes were still visible in the areas covered by dermis. The dorsal and anal fin were not found, while the pectoral, pelvic and caudal fins were visible but, as in all the previous specimens, their structural robustness was lost. No internal organs were identifiable, save for the intestine bulb, which was flat and pale in colour.

### Comparison between the carcasses decaying in sediment (SNDCTRL) and the XCT scanned carcasses

Our experiment investigating the impact of XCT scanning on decay resulted in two datasets for each carcass: one with decay states recorded from the dissections (described in the previous paragraph), and one with decay states recorded from the tomograms. This allowed us to compare the resulting percentages of lost morphological characters and to utilise Spearman’s rank correlation coefficient to test whether XCT scanning impacts the sequence of anatomical character loss of a decaying carcass. Specimen # 39 from the SNDCTRL experiment was used as the comparator.

We first performed the comparison and the analysis using the datasets resulting from dissections. We record advanced decay states for most of the XCT scanned carcasses during dissections (*n* = 5), with extensive disarticulation of the skeleton and most of the internal organs lost. The remaining carcasses (*n* = 3) lost some internal organs (e.g. swim bladders, liver, pancreas) and were largely articulated. As a result, more than half of the carcasses (*n* = 5) have a percentage of lost characters 2-3 times higher than the control, most of them (*n* = 3) are among the carcasses XCT scanned twice during the experiment (Table 1). This indicates that XCT at the start of the experiment may cause an elevated rate of decay when compared to carcasses buried in quartz sand. The Spearman’s rank shows that the majority of carcasses are statistically similar to the control in terms of sequence of anatomical character loss (*n* = 6) (Table 1). The carcasses that deviate from this trend are also the ones with the highest percentages of lost characters (samples A, F) (Table 1; Appendix S1).

**Table 1:**
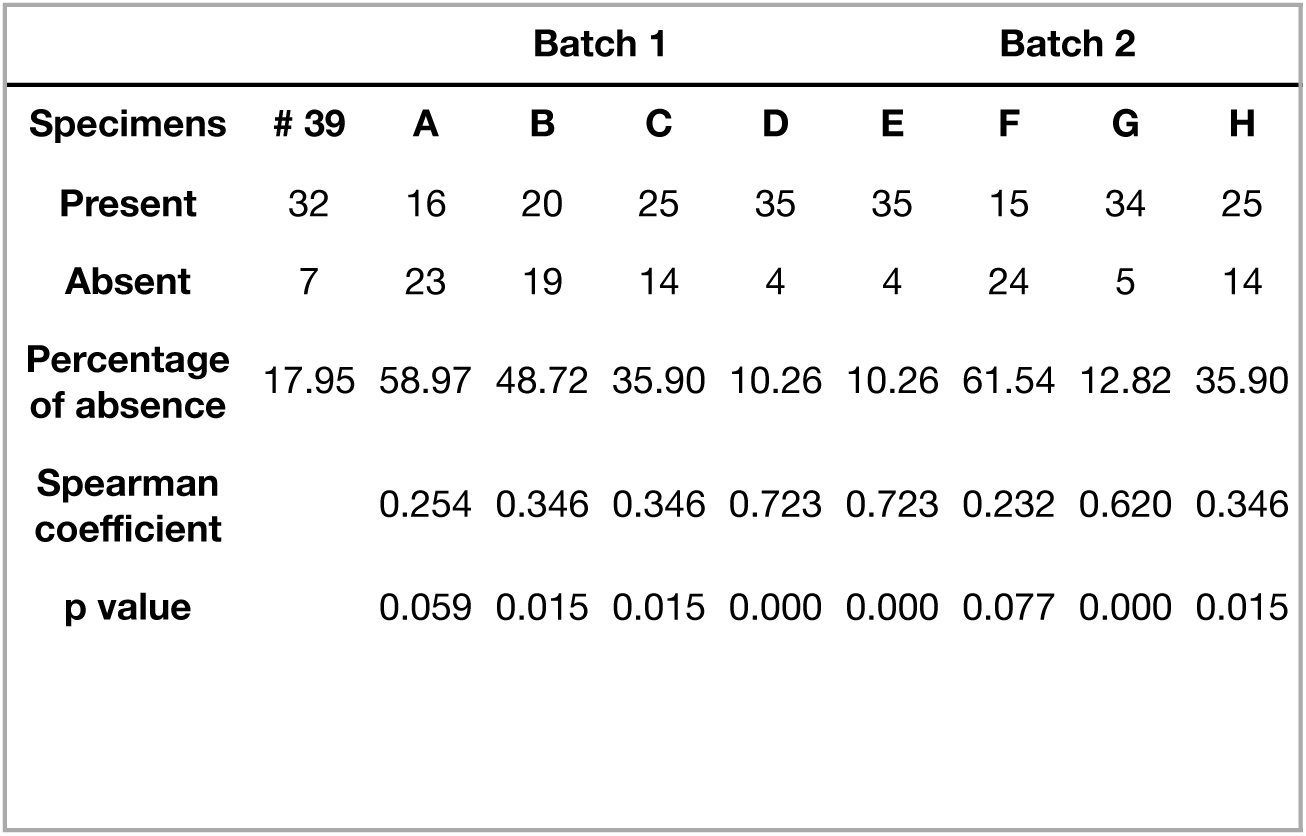
Summary of comparisons and Spearman’s rank analysis using presence/absence data from dissection. **Footnote:** Specimens A-D are in batch 1 (XCT scanned at the initiation of the experiment and at its termination on day 50); specimens E-H are in batch 2 (XCT scanned at the termination of the experiment on day 50). Specimen # 39 from our sediment-based control experiment (SNDCTRL) is the reference for both analyses. P-value significance level: 0.05.

The datasets resulting from the tomograms are largely limited to the mineralised tissues (Supp.Mat.). It is known that XCT scanning does not allow to image soft tissues without staining agents, in fact the non-mineralised morphology of the carcasses appears featureless and grey in our tomograms. Nonetheless, we were able to confirm the presence and full articulation of the skeleton of all the samples (Fig. 3). We combined the dissection and tomographic data for each carcass, reporting any character visible in either method as present. We find that this combined dataset has generally lower percentages of absent characters when compared to the dissection dataset alone, although most carcasses (*n* = 5) still have higher percentages of loss than the control (Table 2). In addition, the Spearman’s rank correlation coefficient show that all carcasses have a constant sequence of anatomical character loss (Table 2; Appendix S1).

**Table 2:**
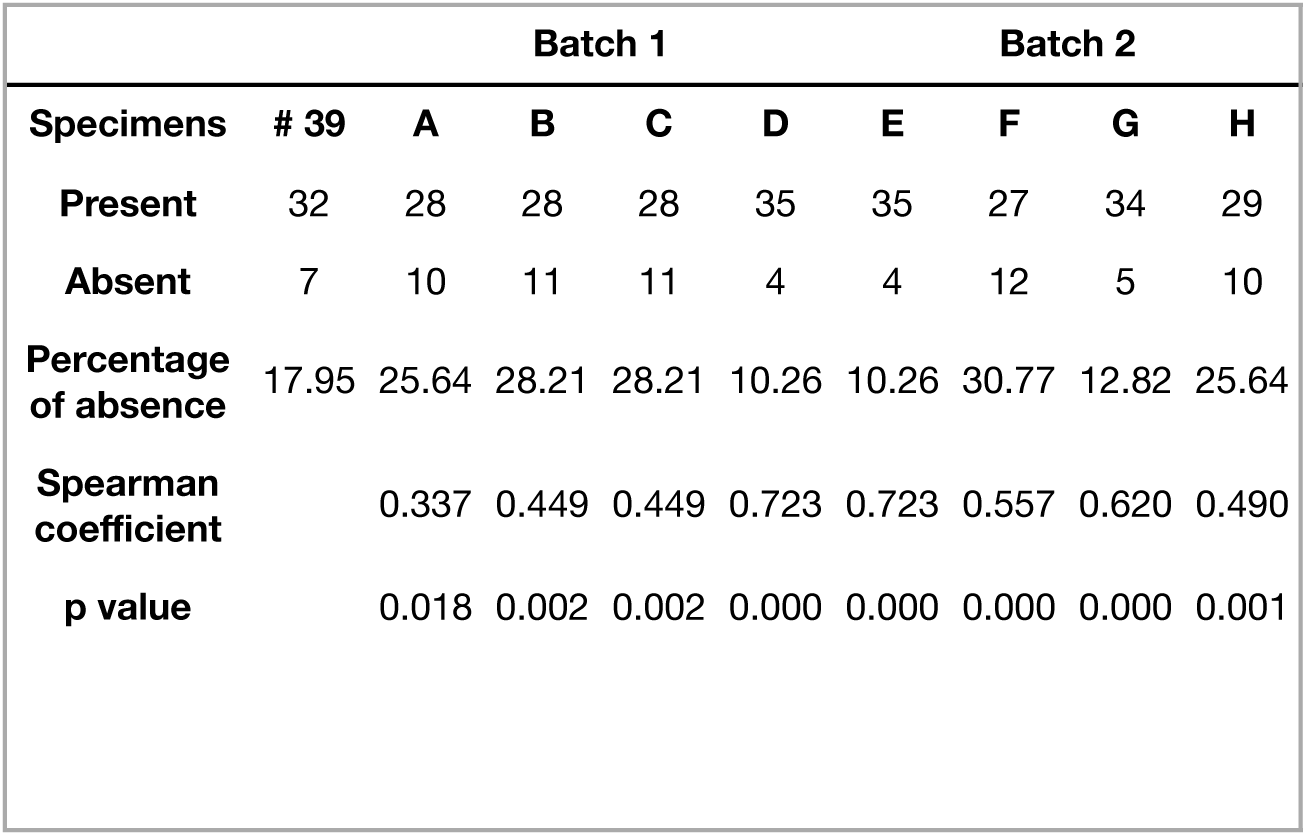
Summary of comparisons and Spearman’s rank analysis using combined presence/ absence data from dissections and tomograms. **Footnote:** Specimens A-D are in batch 1 (XCT scanned at the initiation of the experiment and at its termination on day 50); specimens E-H are in batch 2 (XCT scanned at the termination of the experiment on day 50). Specimen # 39 from our sediment-based control experiment (SNDCTRL) is the reference for both analyses. P-value significance level: 0.05.

### Carcass orientation within substrate and 3D model mesh comparison

Here we report the results of the mesh comparison conducted on specimens A-D, which were XCT scanned on both day 0 and day 50 (Fig. 4).

**Figure 4:**
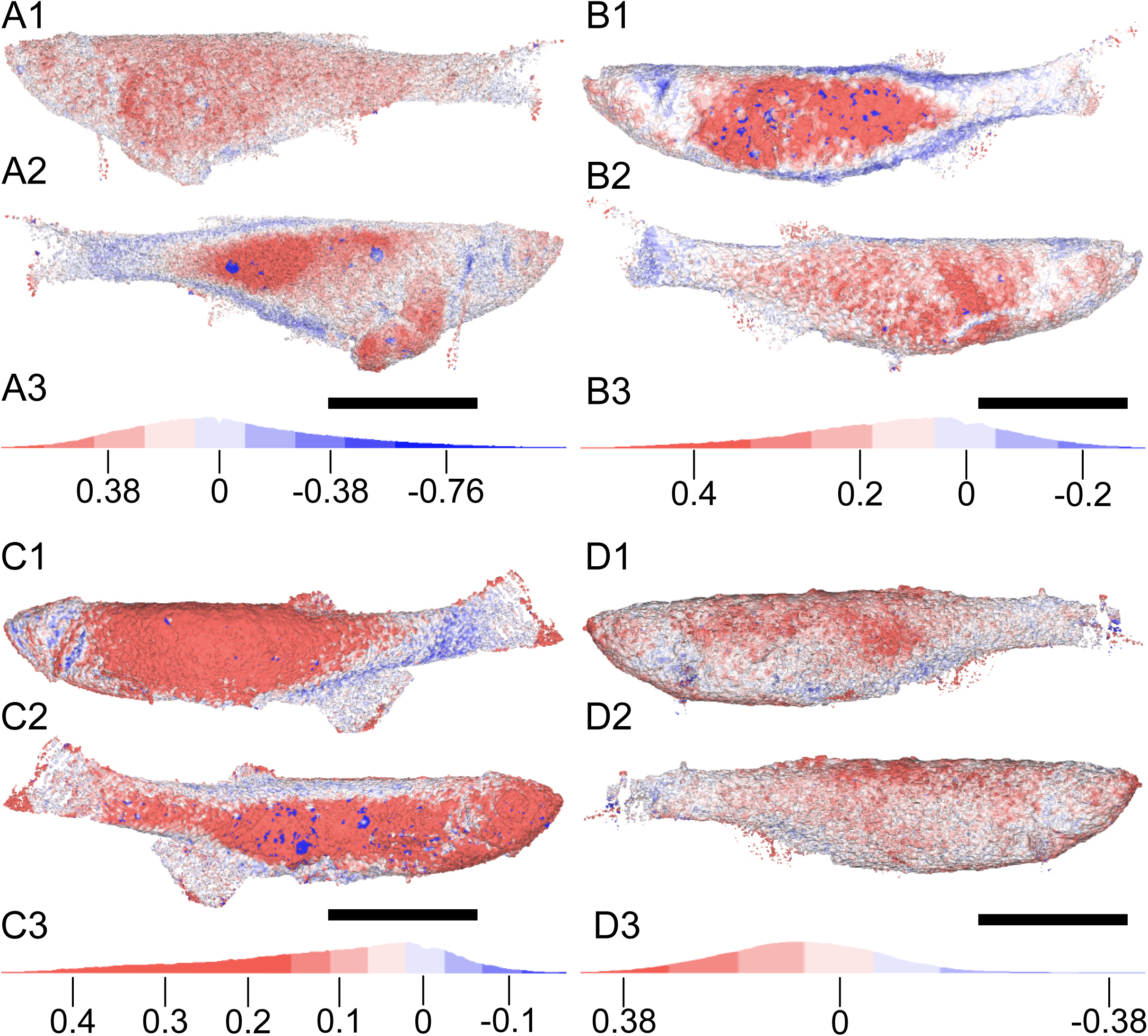
Comparison of carcass volume change during decay. CT renderings of specimens (A-D) at day 0. A1-A2, specimen A. B1-B2, specimen B. C1-C2, specimen C. D1-D2, specimen D. Volume changes of the carcass at day 50 are plotted on the renderings as a heatmap. Red hues indicate areas of volume decrease over time (i.e. day 0 models are larger than day 50 models); blue hues indicate areas of volume increase over time (i.e. day 0 models are smaller than day 50 models). A3, B3, C3, D3: keys showing distortion values in mm and corresponding colour hues. Specimens A - C decayed laterally (A2, B1, and C2 are upward). Specimen D decayed with dorsal edge upward. Scale bar: 1 cm. Mean distance (mm): -0.06063 (specimen A); 0.1066 (specimen B); 0.1204 (specimen C); 0.1393 (specimen D). Standard deviation: 0.3933 (specimen A); 0.1798 (specimen B); 0.1654 (specimen C); 0.1995 (specimen D).

Specimens A and B were oriented on their lateral side at the initiation of the experiment (left lateral for A, right lateral for B). Both carcasses show volume reduction over time on both lateral surfaces. Post-decay, the upward facing side of each fish displays an ovoid area of volume reduction from behind the skull to the caudal peduncle (0.1–0.5 mm) and, in specimen A, an area of volume reduction is also observed in the extruded intestine (Fig. 4A1). Narrow areas along the dorsal and ventral margins of the body, as well as the posterior margin of the skull and operculogular series, show a volume expansion instead (0.2–0.5 mm) (Fig. 4A1, B1). The volume reduction on the downward facing side of the fish is less pronounced but more widespread across the entire lateral surface of the body (Fig. 4A1, B1). A modest volume expansion is seen along the dorsal margin of the specimen and the caudal fin.

Specimen C decayed lying on its left lateral side, as with specimen A, but curved into a wide “U” shape such that the head and caudal fin extended higher up in the sediment than the rest of the body. Areas of equally pronounced volume reduction (> 0.2 mm) are seen on both sides of the body (Fig. 4C1). On the downwards facing side, the reduction in volume is concentrated between the back of the skull and the anal fin, and particularly pronounced on the portion of the body that was the lowest in the sediment. Small areas of expansion are seen along the ventral margin of the body in proximity to the anal fin, the caudal peduncle, and around the skull and operculum. The upwards facing surface of the specimen displays a wide area of volume reduction running the length of the body, from the head to the tail. Very minor expansion is visible along the dorsal margin of the body, as well as in a narrow area near the anal fin (Fig. 4C1).

Specimen D decayed in the sediment with its dorsal margin facing upwards. This orientation caused volume reduction to be concentrated in the dorsal and dorsolateral areas of the body (0.1-0.3 mm), with some minor reductions in the ventral region and around the mouth (Fig. 4D1). Expansion with time is minor (typically < 0.2 mm), and localised in the ventrolateral regions of the body, around the caudal fin, and in correspondence of the operculum (Fig. 4D1).

Our data suggest that the orientation of the carcass has an impact on the amount of decay. When comparing the amount of decay on day 50, carcasses that were buried laterally have higher percentages of character loss (specimens A, B, C; 58.9-35.9%) than the one buried dorsally (specimen D; 10.2%) (Table 1).

## Discussion

Our control experiment confirms that the loss of anatomical characters during decay in zebrafish is not random. To our knowledge, this is the first time this has been demonstrated in a non-cyclostome vertebrate (e.g. Sansom, et al. 2010; Sansom et al. 2013; Murdock et al. 2014; Sansom 2014). This is a taphonomically important finding, as it expands the groups of organisms that have consistent, repeatable, and therefore modellable, decay sequences. This lends further evidence that decay sequences of extant organisms can be used to interpret the fossil remains of related ancient organisms. Furthermore, our secondary experiment shows that burial in a quartz sand substrate does not alter the sequence of anatomical character loss in zebrafish carcasses (Fig. 1; Table S1). This is unsurprising, as previous decay experiments that utilise sediments have not generated results where the carcasses deviate from ‘expected’ decay sequences of anatomical character loss (e.g. Wilson and Butterfield 2014).

In fact, one of the key findings of Wilson and Butterfield (2014) is that the incorporation of sediment impacted the rate of decay; they investigated the impact of sediment type on decay by burying polychaete and shrimp carcasses in kaolinite, calcite, quartz, and Ca-montmorillonite sediments. Their results demonstrate that some mineralogies ‘enhance’ decay, while others can retard it, and that the effects are taxon-specific. They noted that *Nereis* buried in quartz sand appeared to have ‘accelerated’ decay, and our data confirm a similar effect in zebrafish (Fig. 2).

Unfortunately, Wilson and Butterfield (2014) only examined the carcasses at the termination of a 4 month experiment in a qualitative study making it difficult to draw direct comparisons with. Our high-resolution temporal data allows a more detailed investigation of the timing of anatomical character loss in carcasses buried in sediment. One interesting result is that in the first 16 - 18 days of zebrafish carcass decay, we observe a faster onset of loss in the specimens buried in quartz sand (SNDCTRL) than in the control (CTRL) and a faster decay rate (Fig. 1; Fig. S1-S2).

What causes quartz to hasten the onset of decay and increase the rate? We can exclude the effects of any enhanced chemical interactions between the quartz and the carcasses: quartz sand is inert, and we found no evidence of early or enhanced decay of the external surfaces, such as the scales or dermis, in any of the carcasses. In fact, our data show that soft tissue decay proceeds from the internal tissues outwards, with organs lost first, then musculature, and finally (beyond the timescale of our investigation) dermis. It has been shown experimentally that decay by-products and fluids in the proximity of a decaying carcass create a localised geochemical environment that can influence decay rates and soft tissue preservation potential (e.g. Sagemann et al. 1999; Clements et al. 2017; Clements, et al. 2022). Quartz sand has relatively high permeability, thus we would expect this localised decay environment to diffuse away from the carcass; given the lack of any kind of pore water flow in our experiment, it is possible that the sand grains prevent or at least slow the diffusion of fluids away from the proximity of the carcasses, resulting in a closed microenvironment that enhances decay (as per McCoy, et al. 2015). Our data supports the hypothesis that the formation of localised geochemical environments around a carcass in quartz sediment must occur rapidly, more so than in water alone, and this may explain why onset of decay occurs sooner. Wilson and Butterfield (2014) noted that some clays, such as kaolinite, appeared to retard decay, and while they too would act to trap decay fluids, clays have been experimentally shown to inhibit bacteria (McMahon, et al. 2016). Eventually the pronounced rate of decay in quartz sand slows to be more similar to the control; presumably, as decay proceeds, the exhaustion of metabolites within the carcass slows bacterial respiration, causing the intensity of geochemical gradients around the carcass to diminish, slowing the rate of decay which will eventually normalise with the sediment-free control.

### The impact of XCT scanning on carcasses in decay experiments

Conceptually, techniques such as XCT scanning can negate the need for exhumation during decay experiments by non-destructively imaging carcasses *in situ*. However, until now, it was not clear if subjecting a carcass to X-rays can have an adverse effect on decay, potentially invalidating the use of XCT scanning in decay experiments.

Our data suggest that exposure to X-rays from XCT imaging at the initiation of a decay experiment may have an impact on the rate of decay, as the carcasses XCT scanned on day 0 show generally higher amounts of decay at the termination of the experiment, compared to the carcasses XCT scanned only at the termination of the experiment (Table 1). As noted previously, rate of decay can be influenced by environmental variables, especially temperature (Briggs and Kear 1993; Briggs and Kear 1994). Within our experiment, it is difficult to determine whether the increased decay rate was caused directly by the XCT scanning or by temperature variations in the environment that the samples were stored in during the analysis (see Methods section). Furthermore, sample agitation could be a factor, although careful handling and burial in quartz sediment attempted to minimise this as much as was practicable. Further investigation is required to fully resolve this, although we note that an elevated rate of decay does not render XCT as a non-practical tool for decay experiments. Importantly for decay experiments, our results indicate that the sequence of anatomical character loss is not impacted by exposure to XCT scanning, when compared to the control (Table 1-2). As it is believed that the sequence of anatomical character loss is governed primarily by the interaction between bacteria and the intrinsic structural properties of biological tissues within a carcass (as per Briggs 2003; Hilal et al. 2021), we hypothesise that the internal microbial communities do not appear to be affected by exposure to X-rays at the commencement of the experiment. Therefore, our results suggest that XCT scanning is a valid technique for use in decay experiments at any point during an experiment, and it is theoretically possible to XCT scan a carcass multiple times.

There are some shortcomings of the application of XCT in decay experiments. The barriers to imaging internal anatomy or the endoskeleton are a consequence of the scarce radiodensity of soft tissues, which causes them to be transparent to X-rays (Metscher 2009a, b; Mizutani and Suzuki 2012; Gignac et al. 2016). Therefore, the impossibility for unaided XCT scanning to image soft tissues means that dissections remain necessary and the advantages of imaging *in situ* cannot be fully explored. In our experimental design, this is exacerbated by the sand and water surrounding the fish carcasses: both media are relatively dense, and water also typically overlaps in density with organic tissues when XCT scanned, limiting the available scan settings (Pauwels et al. 2013). A high beam energy (kV) is required to ensure sufficient penetration of the sediment but decreases the contrast in the resulting images. Sufficient filtration (in the form of thin sheets of metal applied between the source of the X-rays and the specimen) can provide increased contrast by selectively absorbing lower energy X-rays, but increases noise in the resulting images. Thus, the interrelation of a higher kV and the need for filtration in response directly reduces image contrast and image quality in our experiment (per Zwanenburg et al. 2022). We are currently exploring ways to mitigate this by the use of contrast agents, such as Lugol’s iodine, phosphotungstic acid (PTA) or other media (Rawson et al. 2020). Typically, immersion of soft-tissue samples in a staining medium results in passive absorption in relative amounts, based upon the tissue types within the sample (Metscher 2009a). This process creates artificial density differentiation between tissue types, resulting in greater contrast between them during XCT scanning. This ongoing research will define a viable protocol for the use of contrast agents within taphonomic experiments using sediment.

Despite this shortcoming, the ability to non-destructively image a carcass *in situ* has important implications for the accuracy of decay experiments. Tomograms show that carcasses remained articulated within the sediment before extraction (Fig. 3). This is significant, as five out of eight specimens partially or almost completely disarticulated during exhumation (e.g., specimen A, which from the beginning of the experiment had its body cavity ruptured, with partial extrusion of the intestine: Fig. 3A1-A2 and Fig. 4A1-A2). The remaining three carcasses had a similar appearance. This confirms that the exhumation process can be a source of potential bias, especially to carcasses that are in advanced decay states. Our data show that combining the anatomical data collected from the tomograms with the data collected from exhumation and dissection meant samples were typically more similar to the control experiment in terms of anatomical completeness and amount of decay (Fig. 5; Table 2). Therefore, we argue that a combined approach of XCT scanning and traditional exhumation and dissection techniques would constitute best practice, yielding increasingly accurate anatomical presence/absence data in decay experiments that utilise sediment.

**Figure 5:**
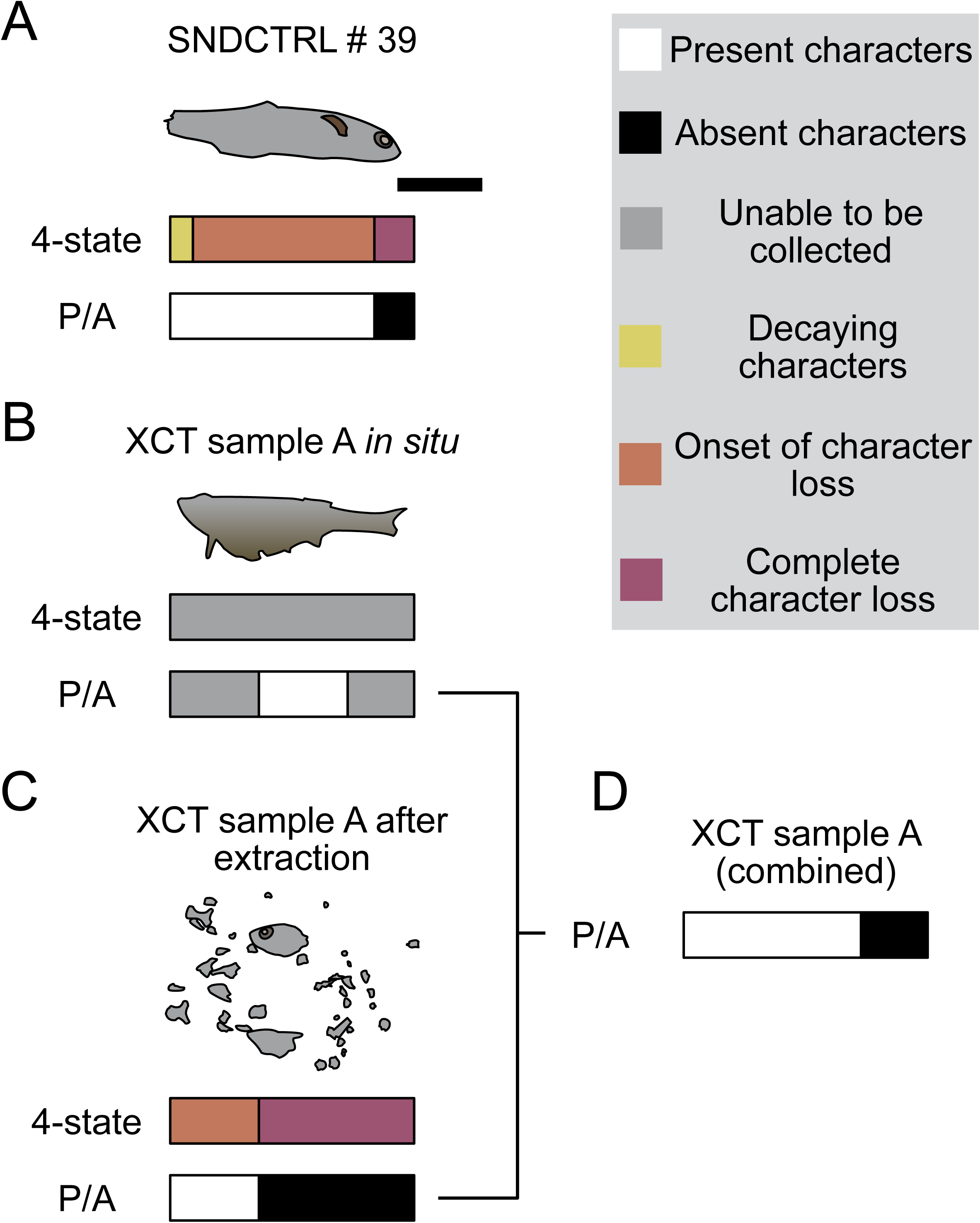
A comparative schematic showing variance in appearance and anatomical character coding of a decaying zebrafish in quartz sand substrate on day 50. A, specimen # 39 from the SNDCTRL experiment after extraction. B, body outline of the 3D model of specimen A from the XCT experiment still *in situ* within quartz substrate. C, specimen A after extraction from substrate. D, presence/absence key for specimen A after combining data from tomogram and dissection. A-C have two keys: (1) ‘4-state’ refers to the percentage of each character coding: pristine (same condition as death), decaying (morphology altered but easily identifiable), onset of loss (morphology altered and difficult to identify), complete loss (no longer observable or recognisable), (2) P/A is a simplified percentage of present versus absent characters. D only has key (2). Scale bar: 1 cm.

### Burial orientation and stabilisation: the role of sediment visualised by XCT

While the primary purpose of this experiment was to investigate the use of XCT in taphonomic investigations, our experiments yield interesting results pertaining to the role of sediment in carcass stabilisation. It is well established that sediment influences carcass articulation during decay: burial prevents transport/agitation by currents, which is a source of disarticulation especially in carcasses that are in advanced states of decay (Duncan et al. 2003; Bath Enright et al. 2017; Bath Enright et al. 2021). Sediment also acts as a substrate that microbial mats can grow onto, thus indirectly enhancing preservation by adhesion to the substrate (the “stick ‘n peel” effect, see Orr et al. 2016) or engulfment of carcasses by the mats themselves, preventing disarticulation (Darroch et al. 2012; Iniesto et al. 2013; Iniesto et al. 2015). In addition, anoxic conditions within the sediment can prevent scavenging organisms from disturbing carcasses, thus promoting the preservation of soft tissues (Allison 1988; Briggs 2003; Clements and Gabbott 2021). Our tomograms show that all the carcasses XCT scanned at the termination of our experiment were fully articulated *in situ* (Fig. 3), but then most of them disarticulated when extracted from the sediment. This confirms that quartz sand had a stabilising effect, maintaining the structural integrity of the carcass for longer periods of time and preventing soft tissue disarticulation, despite the advanced state of decay.

Our experiment also highlights an additional aspect of burial during decay, which as far as we know is unexplored in the literature: the orientation of the carcasses in sediment can influence decay. During the burial phase of our XCT scanning experiment setup we ensured that carcasses were fully buried in sediment, but we did not control their orientation. As a result, orientation varied among the carcasses, and the 3D mesh comparison highlighted a relationship between orientation, patterns of volume variation and, interestingly, amount of decay. Our data show that zebrafish carcasses buried laterally in quartz sand have a distinctive pattern of volume reduction along both flanks, more pronounced on the uppermost flank, and moderate volume increase along the dorsal and ventral margins of the body (Fig. 4A1-C3); this is paired with a higher percentage of lost characters when compared to different orientations (A, B, C from batch 1; F from batch 2; Table 1-2); three of these specimens disarticulated during extraction (A, B, F) demonstrating that the substrate was maintaining carcass integrity. If the carcasses are buried with their dorsal margin uppermost in the sediment, volume reduction is localised mostly dorsally and dorsolaterally, while there is a modest volume increase ventrolaterally (Fig. 4D1-D3). This is accompanied by a lower percentage of lost characters when compared to laterally buried carcasses (D from batch 1; Table 1-2). Specimens E, G, and H were slightly tilted, with one of the dorsolateral margins uppermost in the sediment, and the resulting amount of decay varies (Table 1-2). To our knowledge, this is the first time that the effects of carcass orientation on decay have been observed and recorded in a decay experiment. Our data suggest that carcass orientation could be a currently unknown variable that impacts decay rates, and the entombing sediment may play a role. We hypothesise that the differences in volume variations during decay (Fig. 4) are caused by variations in the orientation of the carcass relative to the application of sediment overburden. If the fish carcass is buried dorso-ventrally, the vertebral column is capable of resisting overburden compression during the early phases of the decay process, causing small volume variations and a diminished amount of decay (Fig. 4D1-D3). Conversely, when overburden loading is lateral (i.e. the carcass is buried on its side), the ribs are not able to resist compaction, so overburden causes larger deformations and higher amounts of soft tissue decay in the same amount of time (Fig. 4A1-C3). In all the specimens but one (specimen G), gas is visible in the tomograms, trapped inside the body (Supp. Mat.), therefore, suggesting that burial orientation does not seem to promote the escape of decay related gases from the carcass.

The effect of burial orientation on carcass geometry when overburdened would probably compound over time and be exacerbated by increasing overburden, compaction, and diagenetic processes. Further investigation is needed.

## Conclusion

As decay experiments increase in complexity, many studies are utilising sediment in their experimental design. However, observation of the carcass is not possible in many ‘natural’ sediments (such as sand, clays, etc.) and so exhumation and extraction are required to collect character loss data, but these processes may cause carcass disarticulation and potential data loss.

In order to mitigate these issues, XCT has been a proposed method to visualise decaying carcasses, however, the impact of exposing a decaying carcass to X-rays is currently unknown. Our data show that while XCT scanning may increase the rate of decay, it does not impact the sequence of anatomical character loss in zebrafish during decay and is therefore a viable technique for non-destructive observation in decay experiments. XCT data confirm that a substrate acts to physically stabilise a decaying carcass, preventing disarticulation over the timescale of our experiment. Interestingly, our data show a new potential variable that affects rate of decay – the post burial orientation of the carcass influences volume change via overburden, and importantly also influences the rate of decay: a carcass lying laterally in sediment decays more than a carcass buried in a dorsal-ventral burial position.

Our control experiments show that burial of zebrafish carcasses in quartz sediment hastens the onset of character loss and increases the rate of decay when compared to carcasses decayed in water alone. Further work is needed to investigate the role of sediment types on this phenomenon.

The use of XCT in decay experiments is limited by difficulty in imaging the internal anatomy of carcasses. However, our data show that XCT scanning during decay experiments allows a more thorough and accurate investigation of a carcass when compared to dissection alone. Furthermore, XCT has other experimental design benefits such as requiring fewer individual carcasses. The work presented here provides a basis for our future investigations seeking to increase the accuracy of XCT scanning in decay experiments (e.g. the implementation of staining protocols).

## Captions

**Table S1: Scores and rankings of characters from CTRL and SNDCTRL.** Characters are in alphabetical order. The scores are calculated for the whole experiment, following the revised methodology from Sansom et al. (2010): pristine = 0; decaying and onset of loss = 0.5; complete loss = 1.

## Acknowledgments

We thank P. Jones and the team from the Husbandry Unit at the School of Biosciences, University of Birmingham, for facilitating access to the zebrafish specimens used in this investigation.

TC was supported by a Leverhulme Early Career Fellowship (ECF-2019-097) during the implementation of this project.

IC was supported by a Central England NERC Training Alliance (CENTA) PhD scholarship via grant NE/S007350/1.

SG was supported by a Royal Society Dorothy Hodgkin Research Fellowship (no. DH160098). The X-Ray Computed Tomography (XCT) data used in this article was acquired using the Free-at-Point-of-Access scheme at the National Facility for X-Ray Computed Tomography (NXCT) and carried out at the Centre for Imaging, Metrology, and Additive Technologies (CiMAT) at the University of Warwick under the EPRSC Project Number (EP/T02593X/1).

## Author Contributions

**Conceptualisation**: TC. **Data curation**: IC, PW, SG. **Formal Analysis**: IC, TC, PW. **Funding Acquisition**: SG, TC for studentship; SG for lab equipment; supplemented by funding application by IC & TC. **Investigation**: IC, TC, PW, AJ, JA. **Methodology**: IC, TC, PW. **Project**: IC, TC. **Resources**: IC, TC, PW. **Supervision**: TC, SG. **Validation**: IC, TC, SG. **Visualisation**: IC, TC, PW. **Writing – original draft**: IC, TC. **Writing – review & editing**: IC, TC, SG, PW, AJ, JA.

## Data archiving statement

Additional supporting information can be found in the online version of this article (doi to be populated). During the review process, supplementary files will be available at the following links: https://www.dropbox.com/scl/fo/3som902fu98ah561aykc7/AOsOCvY0xhKbRIPTFKnEn80?rlkey=dq5p5s0kqb89f80tlbdbcqxvg&dl=0 for the following files:

-Appendix S1: spreadsheet with character matrix for the CTRL and SNDCTRL experiments, datasets for the XCT scanned specimens and legend for character states.

-Figure S1: bar graph comparison between 4 states percentages of the CTRL and SNDCTRL.

-Figure S2: line graphs of the abundance of each character state in the CTRL and SNDCTRL.

-Table S1: scoring and rankings of the Spearman’s rank analysis between characters of the CTRL and SNDCTRL.

The .ply files of the specimens can be found at the following links:

-sample A_day_0: https://www.dropbox.com/scl/fo/qwbqkd0w9895ej6lrqytq/AOybqGghrobmJXxvBh_mnxU?rlkey=tu4phtfkx0sztmx41d60ios8n&dl=0

-sample B_day_0: https://www.dropbox.com/scl/fo/zn2y0whj88qw1ew9yiwes/AJZScu3YOWTEX8SQ-xL0C-U?rlkey=96nfc71n8z3hkuu8b8i3h5xeq&dl=0

-sample C_day_0: https://www.dropbox.com/scl/fo/psycbrop2vqfzh5n56vae/AAgXHQq3mYr_7Nzb-CgYz2o?rlkey=f06g6xdh80eo2v3ukppviv4nw&dl=0

-sample D_day_0: https://www.dropbox.com/scl/fo/ns4v9o2fjyrw8647c75uk/AB6GrdJvAOBdJ--_FX6yG6w?rlkey=t4qch8zhnqszs0qcrcm18ayr1&dl=0

-sample_A_day_50: https://www.dropbox.com/scl/fo/lhnef0djigv46y88emirj/AB3KPSa7Oe43og_Q9M0kwTM?rlkey=tzj1mjvp5y6v2p1ccjjvok2za&dl=0

-sample_B_day_50: https://www.dropbox.com/scl/fo/sr5wxjn5427ttzai8lwgh/AA2gSqP0J-utOHECRC5pfgU?rlkey=3ji6fejv9sakd8cp2svqp837c&dl=0

-sample_C_day_50: https://www.dropbox.com/scl/fo/zn0r8gkfzgd9gr26cgn59/ADdclvutmuV1Bcn6S6a-p8E?rlkey=2ahyvufhplbmace1clh2jdekj&dl=0

-sample_D_day_50: https://www.dropbox.com/scl/fo/9n2m2drpnzf1udgjx24ss/AA0CliYiHSKrE28MyqDMjuc?rlkey=bdpim3lcw87q6oa7n1oi089yb&dl=0

-sample_E: https://www.dropbox.com/scl/fo/jdbipbt982h7vzgal7hht/AAnkf3PieU9qm5zkGGl-yZk?rlkey=uof1xx7nxm1u1taoniiytrxpb&dl=0

-sample _F: https://www.dropbox.com/scl/fo/5mhdmfxymtigtobjf0216/ADxARgjpO4CIzbUgdTa_W_o?rlkey=dcyb7wf2wwp5b5tj2feelbkut&dl=0

-sample_G: https://www.dropbox.com/scl/fo/n1724tsu45kp74jnzosfz/AAiHy1gn-kBjyxcJpUo1_MY?rlkey=ndxx5ek6t9w4rf2cfhw3rlu94&dl=0

-sample _H: https://www.dropbox.com/scl/fo/y9g38sxtiwiqdqb05uszx/AJWak9BhBFW_cPnnGqzhfL4?rlkey=fndyngct6mg09zyjw51jtoiq7&dl=0

The raw scan data can be found at the following links:

-sample A _day _0: https://www.dropbox.com/scl/fo/bxh88iyj6jkp6wl8hfrsw/AKeJT6XVrb58yTnPqHMJl_4?rlkey=u0otvmoj8tw7upuh08xdnz4ma&dl=0

-sample_A_day_50: https://www.dropbox.com/scl/fo/dieni2hz892028gjejwi1/AEIgQBQIdaJtmGsui9Z3Fo8?rlkey=l846mhufeswnxmz4cnzb1d89k&dl=0

-sample B_day_0: https://www.dropbox.com/scl/fo/eg2zyyyohnw7pkp2glaru/AB0yJ0V2L8jhScRG5iiKx1U?rlkey=b9sue7xhhavan454aumb67peb&dl=0

-sample_B_day_50: https://www.dropbox.com/scl/fo/3s97xqygevbqxj7f1piml/AHhi3ve-9-08_WNqg-xeqYw?rlkey=lkcri7ell9b4ugeijm3rmk7pp&dl=0

-sample C_day _0: https://www.dropbox.com/scl/fo/coltjy9efkldiko8evreq/AGjunN9WzNi32IsvQsqdip8?rlkey=0j5apz8bmrrzll8tpfjpo8r0f&dl=0

-sample_C_day_50: https://www.dropbox.com/scl/fo/i9lezryy2jc0ipf3qa2oq/AEA9118AG0rXR4ltFiQ70sU?rlkey=4d0uzb7ywgzyaauzfq3t3v8sw&dl=0

-sample D_day _0: https://www.dropbox.com/scl/fo/1r96irf3o0iixf88iagaz/AMEdcassP_QNpJGsgK7O6sg?rlkey=4zx5nmv9slh1pdjbixa3itr2k&dl=0

-sample_D_day_50: https://www.dropbox.com/scl/fo/gwu82o637aaxpxvd54i43/ABku8nePIWzp6xgahbhcze4?rlkey=083a8eru5rx0yg9wyi45yj28f&dl=0

-specimen_E_day_50: https://www.dropbox.com/scl/fo/8a77flnjse2d1dj9zalvw/ANGejq0Omm3ACkLJjPZvTEw?rlkey=5yzytl84sy3zrv2rtvqx1we1t&dl=0

-specimen_F_day_50: https://www.dropbox.com/scl/fo/7z16mluqdzzqt6gbptgq3/APLX7tWALB8m3aYEey-mJyc?rlkey=z102vt52dxwnn1q27atuzyucz&dl=0

-specimen_G_day_50: https://www.dropbox.com/scl/fo/qy54ur9cgqf0s5i17fzk1/APn8Fo5XvIErMT_jvZ13eEA?rlkey=8mbpe52yzpgugdkh18e73z3l1&dl=0

-specimen_H_day_50: https://www.dropbox.com/scl/fo/00v8dqf61dz7k3w48f850/AA0S-46lwaM2u4e_swGbyQo?rlkey=0omuayr5z9b2kleicrmoiej5m&dl=0

