## Supplementary figures and images for "Investigating the impact of X-rays on decay: X-ray computed tomography as a non-invasive visualisation technique for sediment-based decay experiments"

### figs1_166.tiff

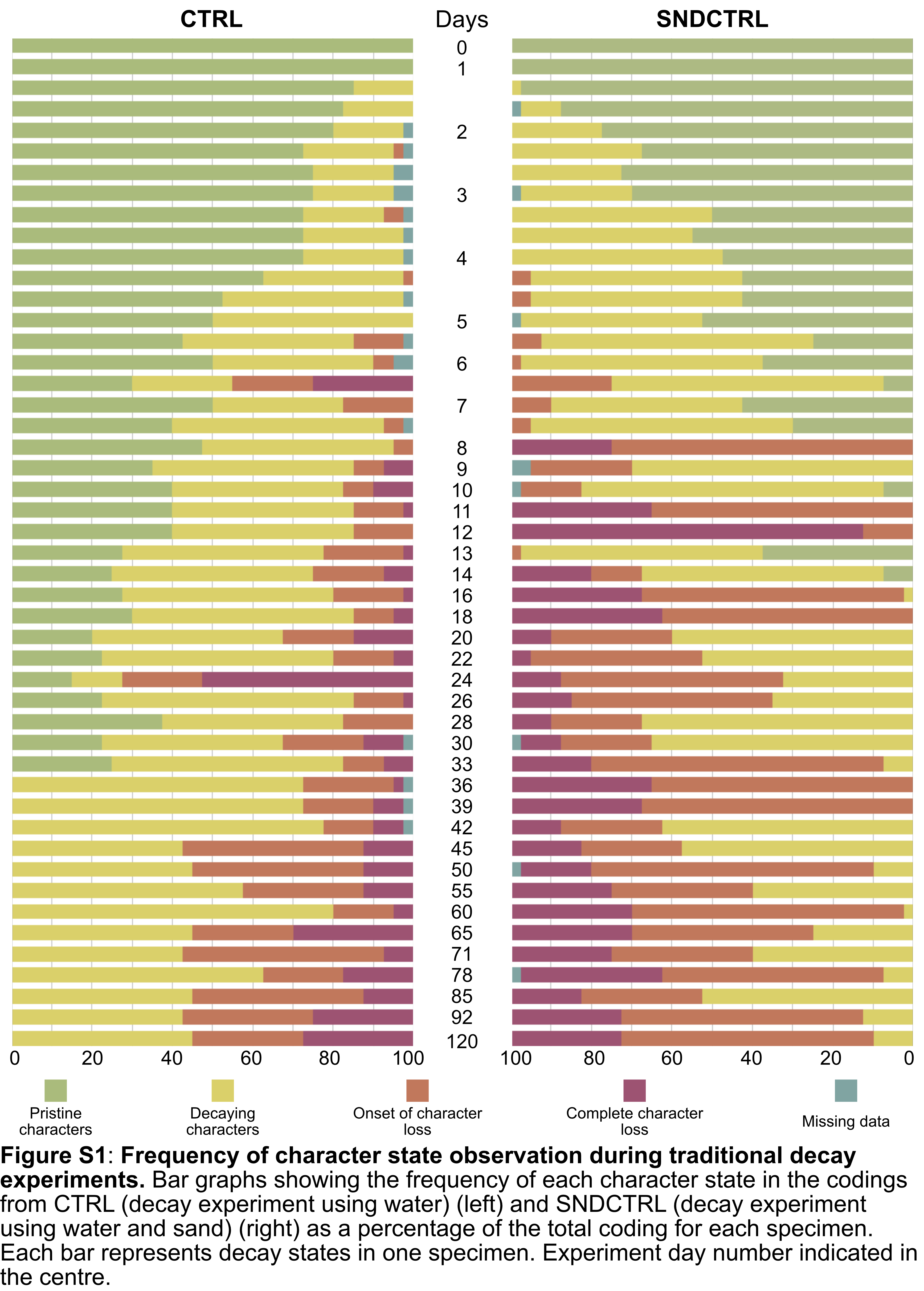

### figs2_166.tiff

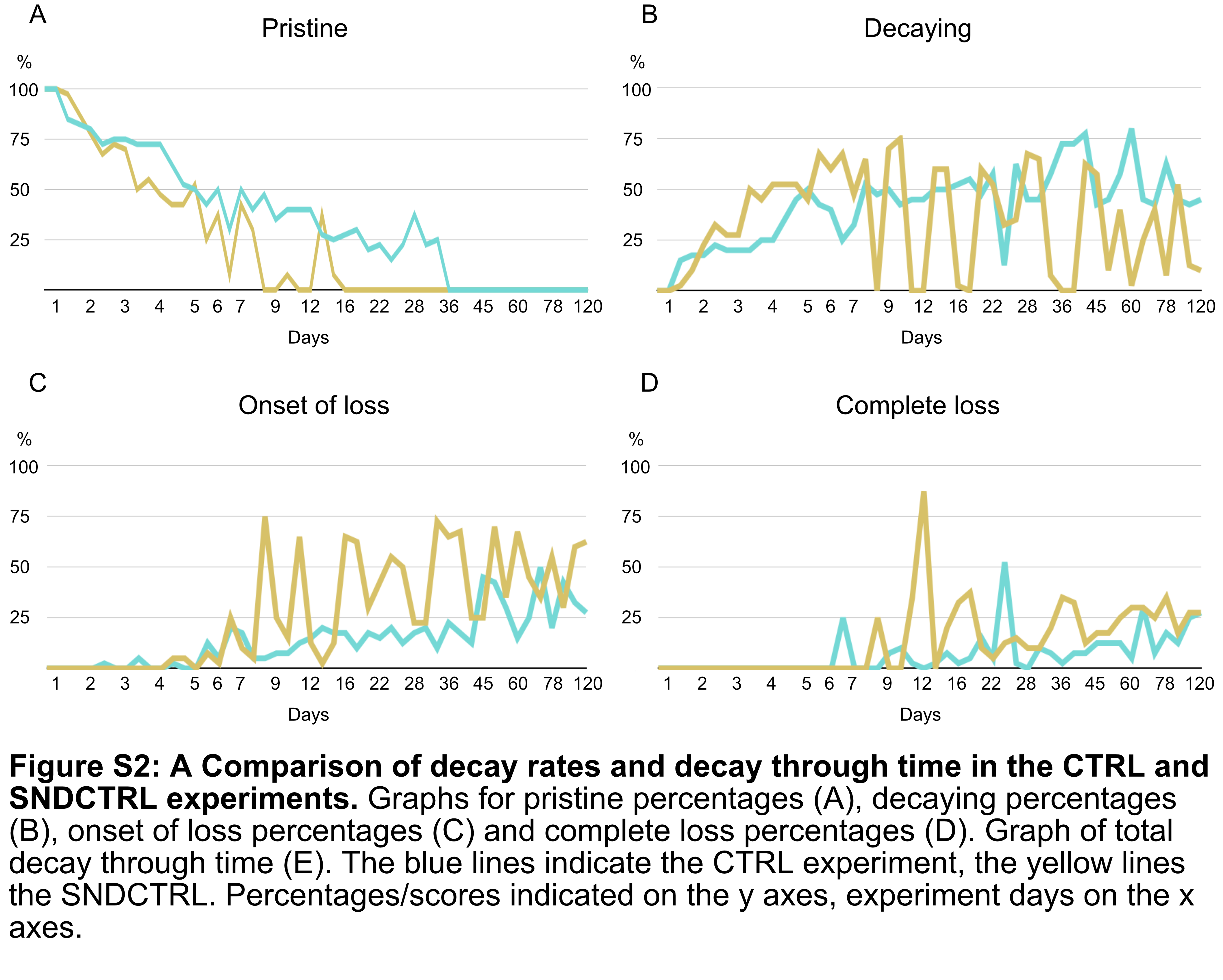
